# Comparative analysis of Wnt signaling-related proteins in normal, benign, malignant and metastasised human liver tumors

**DOI:** 10.1101/2020.11.28.399865

**Authors:** Bushra Kanwal, Yoh Zen, Maesha Deheragoda, Michael Millar, Christopher Thrasivoulou, Sheikh Tamir Rashid, Aamir Ahmed

## Abstract

**Background and Aims:** Liver cancer comprises of benign or malignant tumors including hepatocellular carcinoma (HCC), cholangiocarcinoma (CC), hepatoblastoma (HB), and other rarer tumor types. There is evidence of aberrant Wnt signaling during initiation and progression of HCC, CC and HB.

**Methods:** We investigated the expression of Wnt/β-catenin transcription related proteins, Cyclin D1, c-Myc, Fra-1 and Pygo-1, in human liver tumors by using an unbiased, quantitative immunohistochemical (qIHC) approach.

**Results:** Semi-automated, unbiased quantitation of individual proteins revealed reduced expression of Cyclin D1 and Pygo-1 in CC (P < 0.0001 and P < 0.01, respectively) and HB (P < 0.05 and P < 0.01, respectively) compared to normal liver (NL). Receiver operating characteristic curves showed Cyclin D1 as a putative marker for CC (AUC > 0.8) that discriminates CC from both NL and HCC (P ≤ 0.0001), and Pygo-1 (AUC > 0.7) as a marker for both CC and HCC (P < 0.01) compared to NL. Combining Cyclin D1/Pygo-1 and applying a logistic regression model further improved the diagnostic potential (classifying 84% of NL and CC cases, P < 0.0001). Quantitative co-localisation of tissue samples simultaneously labeled with the four biomarkers, indicated that co-localisation of both Pygo-1/Fra-1 and c-Myc/Fra-1 was also significantly changed in CC and HCC (P < 0.0001) vs NL. Additionally, co-localisation of Pygo-1/Fra-1, in particular, could also distinguish CC from HCC (P < 0.05).

**Conclusion:** Our results indicate that measurement of Wnt signaling markers could be used to stratify liver cancer.

## Introduction

Liver cancer is the second leading cause of premature mortality, sixth leading cancer type and the third most common cause of cancer-associated deaths worldwide. (1, 2) Liver cancer is categorised as primary liver cancer (PLC) or secondary liver cancer (SLC), based on the site of cancer origin. Hepatocellular carcinoma (HCC) is the most common type of PLC, accounting for 75-85% of all liver cancers cases worldwide, followed by cholangiocarcinoma (CC) (10-15%). (1) Anatomically, CC is categorised as intrahepatic (iCC) and extrahepatic (eCC) based on its location with respect to hilum on the biliary tree. iCC arise above the hilar junction of bile ducts, while eCC arise from within/below the hilum and is sub-categorised as hilar/perihilar (pCC) or distal (dCC) tumour with respect to the location of the cystic duct. (3) Most of CCs (50-60%) are pCCs, while dCC and iCC account for 20-30% and 20% of all CC cases respectively. (3) Hepatocellular cholangiocarcinoma (HCC-CC) and hepatoblastoma (HB) are rare PLCs. (4) HCC-CC has a poor prognosis, and shows clinicopathological characteristics of both HCC and CC. (4) HB is a paediatric tumor that accounts for 80% of all malignant liver cancers diagnosed in children, (5) and is characterised by the presence of incompletely differentiated hepatocyte progenitor cells. (6) Benign liver neoplasms are rare and include focal nodular hyperplasia (FNH) and hepatocellular adenoma (HCA). (7) FNH and HCA are usually asymptomatic and grow slowly with normal liver function tests. (8) SLC is metastasised primary cancer of origin other than liver; pancreatic adenocarcinoma liver metastasis (PAC-LM) and colorectal carcinoma liver metastasis (CRC-LM) are most common SLCs with ~50% of primary cancer patients developing liver metastases. (9, 10) The heterogeneity of liver cancer types is an impediment for diagnosis and prognosis and there is a need to identify specific biomarkers for liver cancer subtypes in order to facilitate diagnosis and prognosis.

Wnt signaling pathway is known to play a key role in liver development, homeostasis and pathogenesis. (11) Wnt signalling is activated by the release of free intracellular calcium and desequestration of the transcription factor/co-activator protein, β-catenin. (12) β-catenin enters the nucleus where it forms a protein complex with other transcription factors (from TCF/LEF family), docking proteins (Bcl9 and Bcl9L from the Legless family), and the coactivators (Pygo-1 and Pygo-2 from the Pygo-family and others). (13) An important endpoint of Wnt-mediated β-catenin stabilisation and translocation into the nucleus is the control of gene transcription via the nuclear TCF/LEF/β-catenin/Legless/Pygo complex; (14) a number of these genes are also transcription factors and include, c-Myc, (15) Cyclin D1, (16) and Fra-1. (17) Here we investigated the expression of four Wnt transcriptional targets, Cyclin D1, c-Myc, Fra-1 and Pygo-1, across different types of human liver tumors using an unbiased, qIHC approach (18, 19) in a large number of tissue samples on tissue arrays (TAs). This is the first report investigating a number of downstream Wnt signaling transcriptional targets in a comparative analysis of liver tumors using qIHC and may provide insights into the mechanisms involved in specific liver tumors and disease outcome.

## Materials and Methods

### Ethics approval, tissue acquisition and microarray construction

Human liver tissue array (HuLiv-TA) was constructed (REC no: 08/H0311/201) using archived formalin-fixed paraffin embedded (FFPE) liver biopsies from 165 patients from King’s College Hospital, London and anonymized. Details of the tissue are given in Supporting Table S1 and Fig. S1. The HuLiv-TA design consisted of randomisation of tissue samples: 15 each from normal liver (NL, tissue adjacent to CRC-LM), pCC, iCC, poorly differentiated HCC (pHCC), well-differentiated HCC (wHCC), HCC-CC, HB, FNH, HCA, PAC-LM and CRC-LM. The experimenters were blinded to the identity or diagnosis of each tissue core on the HuLiv-TA.

### Immunohistochemical staining of the HuLiv-TA

Immunostaining was performed using BOND™ automated staining system (Leica Biosystems, Milton Keynes, UK) and BOND Polymer Refine Detection kit (DS9800, Leica Biosystems, Newcastle, UK) according to manufacturer’s recommendations, as described elsewhere. (18, 19) The list of the antibodies used and the optimisation of antibody staining process are given in Supporting Table S4 and Fig. S2. All HuLiv-TA slides were stained simultaneously under identical conditions, for standardized comparative analysis.

### Imaging and protein expression signal quantitation

Single Ab-labelled DAB-stained slides were scanned at 40x magnification (226 nm/pixel resolution) using a Nanozoomer slide scanner (Hamamatsu Photonics UK Ltd, Welwyn Garden City, UK) and the images of individual cores were extracted using a programmed script. (18) The unbiased quantitative method using ImageJ, described previously, (18, 19) was incorporated into a Python script to execute batch analysis. The script sequentially quantified the amount of tissue and the amount of DAB signal expressed per each core image, based on HSV (hue, saturation, value) model. The expression level for each tissue core was calculated as a percentage area fraction per amount of tissue in pixels and the values were subsequently converted into normal distribution using the probit model. (19) Multiplex IF-stained, HuLiv-TA slides were imaged at 20x magnification using Zeiss Axioscan.Z1 digital slide scanner (Carl Zeiss Microscopy, New York, USA), under identical image acquisition settings. Some tissue cores for all disease types were also imaged using Leica SP8 multiphoton confocal microscope (Leica Microsystems) with a 25x objective (NA 0.75).

### Quantitative co-localisation of protein expression in liver TAs

We have previously shown that not only changes in expression of a protein but also alterations in the co-localisation of two proteins may serve as a biomarker of disease type. (18, 19) To measure the proximity of two proteins, we used a quantitative co-localisation approach described earlier. (18, 19) For the measurement of co-localisation coefficients of target proteins, high magnification IF images from up to 3 different areas of randomised tissue cores (5 – 6 cores each for NL, CC and HCC) were acquired on Leica SP8; this was done using a 63x immersion-oil lens (NA 1.4) with 6x optical zoom and a z-step size of 0.17 μm at 1024 x1024 pixels to ensure at least 3 times Nyquist criteria for subsequent deconvolution.

Resulting Leica image files (*.LIF) were imported into the Huygens Professional software (Scientific Volume Imaging) for the image deconvolution to maximise x, y and z resolution. Deconvolved images were saved as Huygens specific files (*.HDF5) for each fluorophore channel. The HDF5 files were imported into the Huygens co-localisation module and background signal was subtracted using the ‘Gaussian minimum’ method; up to three regions of interest (ROIs) per tissue core (18 ROIs per tissue type) were analysed to calculate Pearson correlation coefficient for co-localisation.

### Statistical analysis

D’Agostino-Pearson normality test and Mann-Whitney U test were performed using GraphPad Prism, version 8.0.0 (GraphPad Software, San Diego, California USA). Protein expression data was plotted as normal QQ plots (for normality), box plots (to demonstrate the range of expression levels), mountain plots (to measure AUC) and ROC curves (to evaluate the sensitivity and specificity of proteins as putative biomarkers for cancer). Probability percentage for the disease outcome of CC and HCC was calculated via binary logistic regression model for all possible combinations of proteins using MedCalc version 15.0 (MedCalc software, Ostend, Belgium).

## Results

### Quantitative analysis of Wnt signaling target protein expression in human liver tumors

Protein expression was analysed in 165 human liver samples, annotated by expert histopathologists (Supporting Table S1). All target proteins included in this study (Cyclin D1, c-Myc, Fra-1 and Pygo-1) were expressed in both normal and diseased human liver tissues (Fig. 1 and Supporting Figs S3-S13). Quantitative analysis of DAB-stained HuLiv-TA slides for all 4 proteins showed significant differences between normal and some liver tumor types (Fig. 2). A reduced expression of Cyclin D1 (P < 0.001) and Fra-1 (P < 0.01) in pCC vs NL was observed; while in iCC vs NL, a reduced expression of 3 target proteins (Cyclin D1, c-Myc and Pygo-1) was observed (P < 0.001, P < 0.01 and P < 0.001 respectively). In pHCC vs NL, the expression of Pygo-1 was reduced (P < 0.001), whereas the expression of Fra-1 was increased (P < 0.01). Pygo-1 and Cyclin D1 showed reduced expression in HB vs NL (P < 0.01 and P < 0.05 respectively). In benign liver tumors, expression of Pygo-1 was reduced, only, in FNH vs NL (P < 0.01); while in SLCs, expression of Cyclin D1 and Pygo-1 was reduced in PAC-LM (P < 0.01) and CRC-LM (P < 0.05) respectively, vs NL. This analysis suggests that in a number of different types of liver tumors, expression of most known Wnt targets, except Fra-1 in pHCC, was decreased compared to NL.

**Fig. 1.**
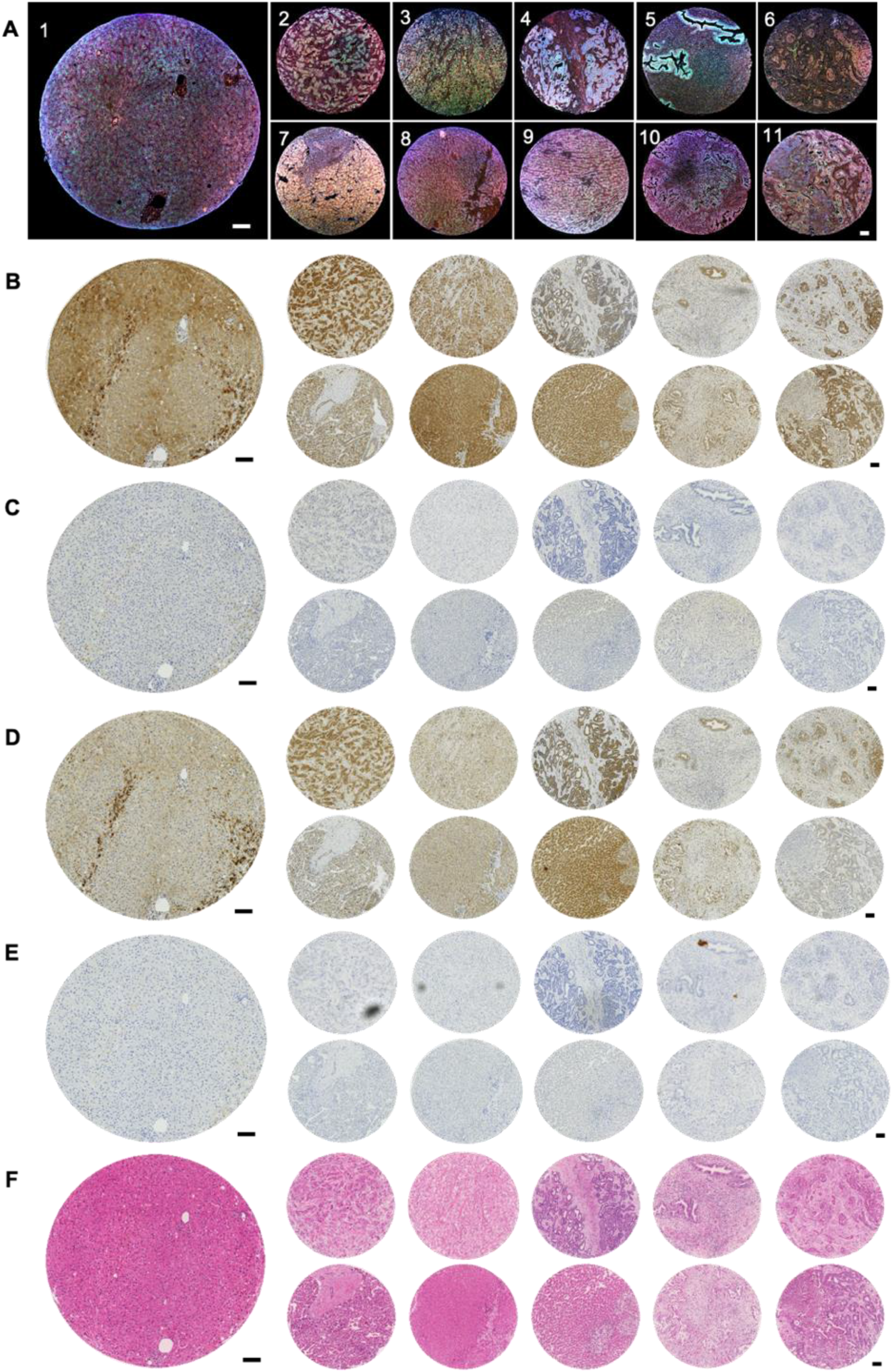
Representative images for various diseased liver tissue types on the HuLiv-TA. (A) Representative multiplex IF images for all tissue types (NL, pHCC, wHCC, iCC, pCC, HCC-CC, HB, FNH, HCA, CRC-LM and PAC-LM numbered from 1 – 11 respectively). Images were acquired on a Leica SP8 at 25x magnification. The composite images represent the co-expression of CyclinD1 (Red), c-Myc (Green), Fra-1 (Blue) and Pygo-1 (Grey). Corresponding DAB-stained images are shown as panel B (Cyclin D1), C (c-Myc), D (Fra-1) and E (Pygo-1). DAB-stained images were extracted from tissue array slide scans obtained from Nanozoomer at 40x magnification. Batch analysis was run on the extracted images to quantify the expression of target proteins (see Methods). Panel F shows corresponding H&E images. Scale bar = 100 μm

**Fig. 2.**
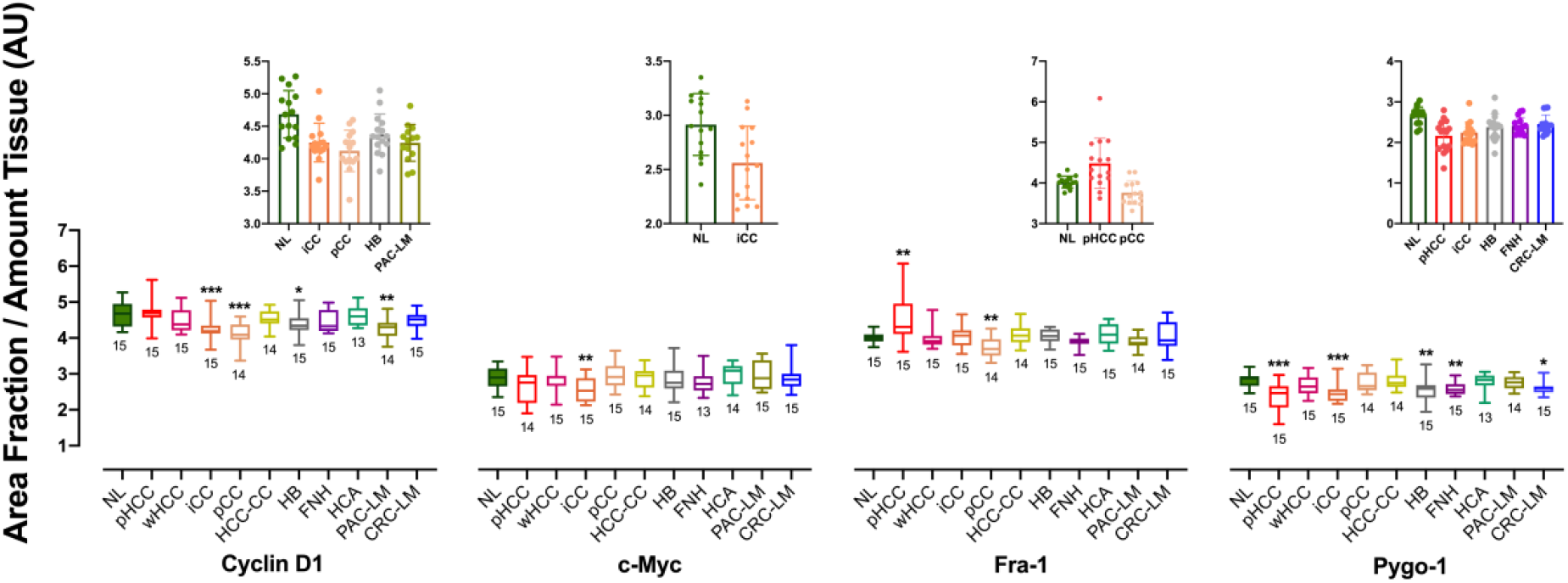
Quantification of DAB intensity to determine expression of Cyclin D1, c-Myc, Fra-1 and Pygo-1 in normal vs benign, malignant and metastasised human liver tissues. Protein expression was calculated as area fraction per amount of tissue in pixels for all tissue cores on the array and transformed into probit values, that were used to create box and whisker plots (AU = Arbitrary units). Significance of difference was calculated between normal liver vs each tumour type using Mann-Whitney U test (* P < 0.05; ** P < 0.01; and *** P < 0.001). The upper panel shows the individual values for significantly different categories plotted as scatterplots with boxes. Sample size (n) for each category is shown below each box.

### Putative diagnostic potential of Cyclin D1, Fra-1 and Pygo-1 protein expression in CC and HCC

Due to the nature of investigation with some liver cancer subtypes that are rarer than others, the subtypes of CC (iCC and pCC) and HCC (pHCC and wHCC) were combined to buttress the statistical sample size for the qIHC analysis. (18, 19) The differential expression was calculated as fold change for each type compared to NL (Supporting Table S2). The mountain plots with a single bin each for NL, CC and HCC showed the differential distribution pattern for all 4 target proteins (Fig. 3). Area under curve (AUC) showed significantly reduced expression of Cyclin D1in CC vs NL and HCC; Fra-1 in CC vs HCC; and Pygo-1 in CC and HCC vs NL, while c-Myc did not show significant difference of expression between NL, CC or HCC (Fig. 3).

**Fig. 3.**
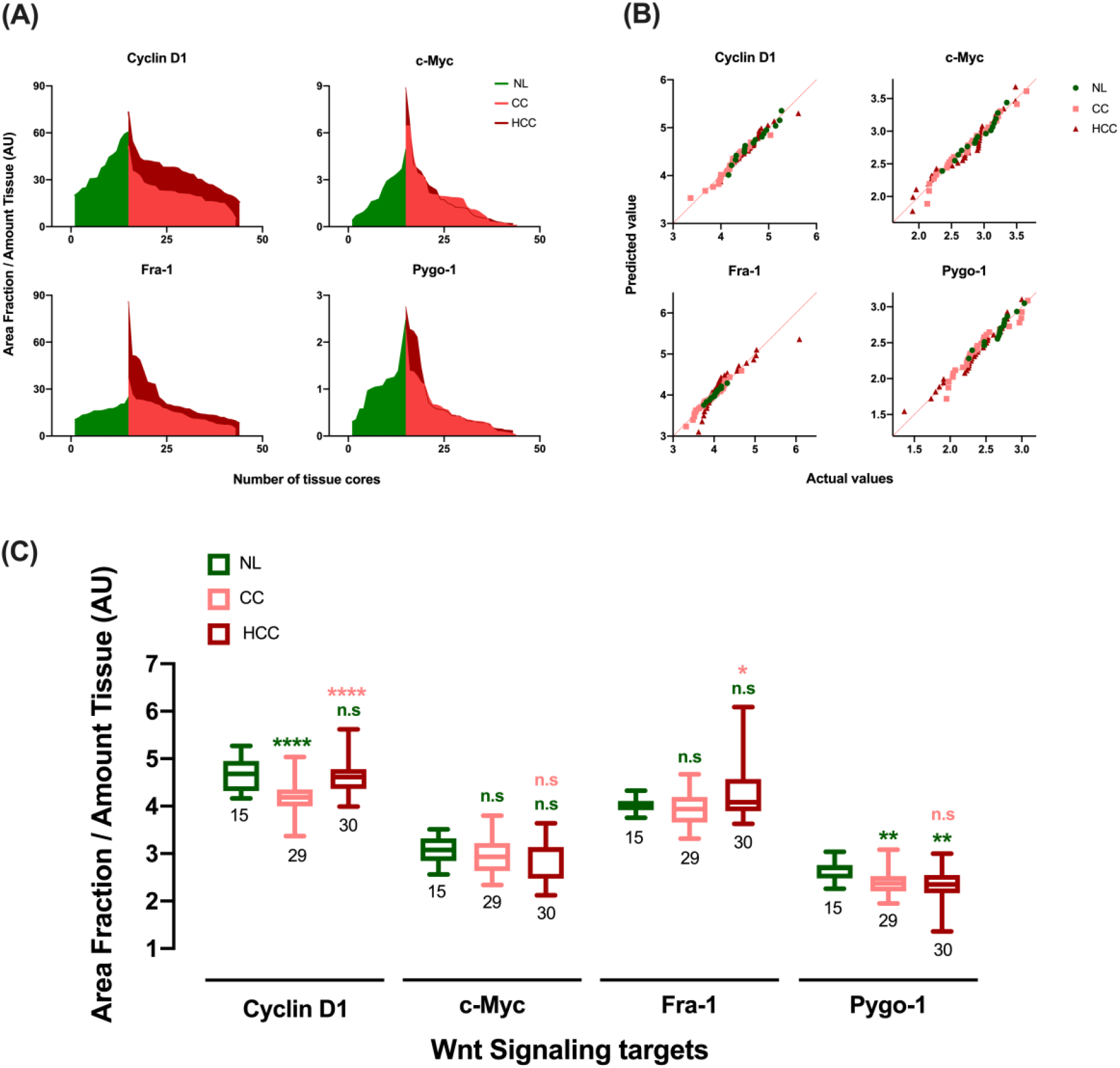
Quantification of DAB intensity for the expression of Cyclin D1, c-Myc, Fra-1 and Pygo-1 in the normal liver, cholangiocarcinoma and hepatocellular carcinoma. Protein expression values for the combined subtypes of CC (iCC and pCC) and HCC (pHCC and wHCC); data is extracted from that presented in Fig. 2. The area fraction per amount of tissue was calculated for all cores as shown in (A). Significant difference between AUC values was seen for NL vs CC and CC vs HCC for Cyclin D1; CC vs HCC for Fra-1; and NL vs CC and NL vs HCC for Pygo-1. The percentage data was then transformed into probit values and Normal QQ plots were created (B). Normalised data was used to create Box and whisker plots (C). Significance of difference was calculated using Mann-Whitney U test for NL vs CC and NL vs HCC, as shown in green above the respective boxes; and for CC vs HCC, as shown in pink above the HCC box (* P < 0.05; ** P < 0.01; **** P < 0.0001; and n.s = not significant). Sample size (n) for each category is shown below each box.

Receiver Operating Characteristic (ROC) curves for the 3 binary conditions; NL vs CC, NL vs HCC and CC vs HCC, were made by plotting sensitivity% (true positive rate) against 100% - specificity% (false positive rate) to illustrate the diagnostic potential of all 4 target proteins for binary classifier conditions (Fig. 4). Discrimination cut-off of 0.5 was used for AUC values of ROC curves implying no predictive power at/below 0.5 and more predictive power (high sensitivity and specificity) at values closer to 1 (Supporting Table S3). Cyclin D1 showed maximum distinguishing potential (i.e. the conditional probability of correctly identifying the two conditions) for NL vs CC (AUC = 0.85, P = 0.0001) and CC vs HCC (AUC = 0.83; P < 0.0001), while Pygo-1 showed significant potential for distinguishing CC and HCC from NL (AUC = 0.76 and 0.77 respectively; P < 0.01). c-Myc did not show any significant distinguishing potential for the 3 binary conditions, while Fra-1 showed significant potential for distinguishing CC from HCC (AUC = 0.67; P < 0.05). The results suggest that Cyclin D1 could serve as a putative diagnostic biomarker for CC, with AUC > 0.8 and likelihood ratios (LRs) ranging from 1.07 – 8.28 and 1.03 – 13.53 for distinguishing CC from NL and HCC, respectively. LRs > 1 indicate that the protein expression value is associated with the presence of disease.

**Fig. 4.**
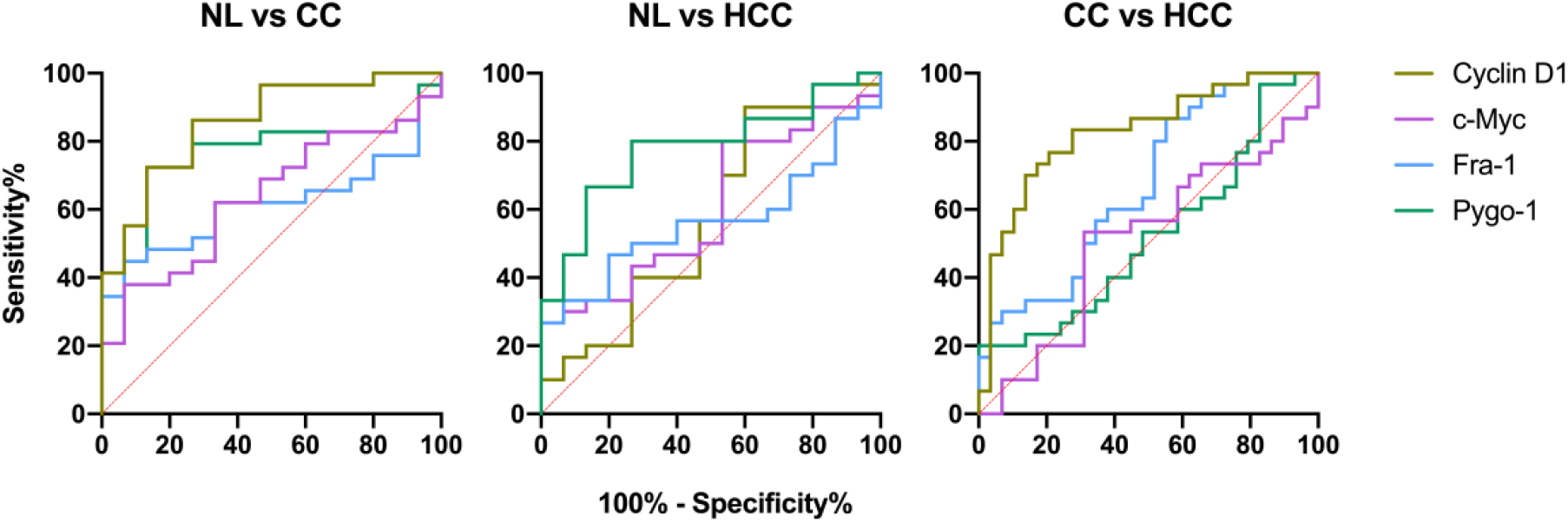
ROC of Wnt signaling target proteins as potential biomarkers of CC and HCC. The ROC curves imply the distinguishing potential of Cyclin D1 and Pygo-1 for CC; Pygo-1 for HCC; and Cyclin D1 and Fra-1 for CC vs HCC. ROC analysis was performed using probit values for the area fraction per amount of tissue for all the cores. The dotted red line represents an AUC of 0.5 with no difference between the respective categories (e.g., NL vs CC; NL vs HCC; and CC vs HCC). The operating characteristic values are given in Supporting Table S3.

The diagnostic potential of the target proteins tested for CC and HCC was improved by incorporating data for all possible combinations of the protein targets into the logistic regression model (Table 1). Cyclin D1/Pygo-1 in combination, predicted the maximum number (84%) of NL and CC cases correctly (P < 0.0001). For HCC vs NL, the greatest number of cases (80%) were classified correctly by combining all four Wnt target proteins (P < 0.01). For discriminating CC from HCC, 5 out of 11 combinations of proteins (Cyclin D1/c-Myc, Cyclin D1/c-Myc/Fra-1, Cyclin D1/c-Myc/Pygo-1, Cyclin D1/Fra-1/Pygo-1 and Cyclin D1/c-Myc/Fra-1/Pygo-1) correctly categorised the highest percentage (77%) of the cases included in this cohort (P ≤ 0.0001).

**Table 1.** Binary Logistic Regression Model to identify the set of biomarkers with improved diagnostic potential for CC and HCC.

| Combinations of Wnt Signaling markers | AUC | Standard error | 95% CI | Significance | Correctly classified cases |
| --- | --- | --- | --- | --- | --- |
| **Normal Liver vs Cholangiocarcinoma** |  |  |  |  |  |
| Cyclin D1 / c-Myc | 0.857 | 0.0581 | 0.719 to 0.944 | P = 0.0001 | 79.55% |
| Cyclin D1 / Fra-1 | 0.860 | 0.0666 | 0.722 to 0.946 | P = 0.0001 | 81.82% |
| Cyclin D1 / Pygo-1 | 0.922 | 0.0394 | 0.800 to 0.981 | **P < 0.0001** | **84.09%** |
| c-Myc / Fra-1 | 0.724 | 0.0753 | 0.569 to 0.848 | P = 0.1250 | 63.64% |
| c-Myc / Pygo-1 | 0.754 | 0.0754 | 0.601 to 0.871 | P = 0.0221 | 65.91% |
| Fra-1 / Pygo-1 | 0.869 | 0.0556 | 0.733 to 0.952 | P = 0.0001 | 81.82% |
| Cyclin D1 / c-Myc / Fra-1 | 0.851 | 0.0692 | 0.711 to 0.940 | P = 0.0003 | 81.82% |
| Cyclin D1 / c-Myc / Pygo-1 | 0.926 | 0.0378 | 0.806 to 0.983 | **P < 0.0001** | 79.55% |
| Cyclin D1 / Fra-1 / Pygo-1 | 0.920 | 0.0409 | 0.797 to 0.980 | **P < 0.0001** | 83.36% |
| c-Myc / Fra-1 / Pygo-1 | 0.871 | 0.0553 | 0.736 to 0.953 | P = 0.0003 | 81.82% |
| Cyclin D1 / c-Myc / Fra-1 / Pygo-1 | 0.933 | 0.0357 | 0.815 to 0.986 | **P < 0.0001** | 81.82% |
| **Normal Liver vs Hepatocellular carcinoma** |  |  |  |  |  |
| Cyclin D1 / c-Myc | 0.647 | 0.0880 | 0.490 to 0.783 | P = 0.2478 | 66.67% |
| Cyclin D1 / Fra-1 | 0.711 | 0.0825 | 0.557 to 0.836 | P = 0.0799 | 71.11% |
| Cyclin D1 / Pygo-1 | 0.813 | 0.0666 | 0.669 to 0.914 | P = 0.0013 | 77.78% |
| c-Myc / Fra-1 | 0.644 | 0.0808 | 0.488 to 0.781 | P = 0.1615 | 62.22% |
| c-Myc / Pygo-1 | 0.813 | 0.0631 | 0.669 to 0.914 | P = 0.0022 | 75.56% |
| Fra-1 / Pygo-1 | 0.782 | 0.0687 | 0.634 to 0.891 | P = 0.0097 | 71.11% |
| Cyclin D1 / c-Myc / Fra-1 | 0.704 | 0.0833 | 0.550 to 0.831 | P = 0.1556 | 71.11% |
| Cyclin D1 / c-Myc / Pygo-1 | 0.847 | 0.0618 | 0.708 to 0.937 | P = 0.0017 | 77.78% |
| Cyclin D1 / Fra-1 / Pygo-1 | 0.813 | 0.0666 | 0.669 to 0.914 | P = 0.0036 | 75.56% |
| c-Myc / Fra-1 / Pygo-1 | 0.813 | 0.0637 | 0.669 to 0.914 | P = 0.0064 | 73.33% |
| Cyclin D1 / c-Myc / Fra-1 / Pygo-1 | 0.844 | 0.0597 | 0.705 to 0.935 | P = 0.0035 | **80.00%** |
| **Cholangiocarcinoma vs Hepatocellular carcinoma** |  |  |  |  |  |
| Cyclin D1 / c-Myc | 0.824 | 0.0551 | 0.703 to 0.911 | **P < 0.0001** | **77.97%** |
| Cyclin D1 / Fra-1 | 0.831 | 0.0538 | 0.711 to 0.916 | **P < 0.0001** | 76.27% |
| Cyclin D1 / Pygo-1 | 0.845 | 0.0522 | 0.727 to 0.926 | **P < 0.0001** | 76.27% |
| c-Myc / Fra-1 | 0.703 | 0.0673 | 0.570 to 0.815 | P = 0.0109 | 59.32% |
| c-Myc / Pygo-1 | 0.539 | 0.0759 | 0.404 to 0.670 | P = 0.6543 | 54.24% |
| Fra-1 / Pygo-1 | 0.718 | 0.0659 | 0.586 to 0.828 | P = 0.0069 | 67.80% |
| Cyclin D1 / c-Myc / Fra-1 | 0.830 | 0.0540 | 0.710 to 0.915 | **P = 0.0001** | **77.97%** |
| Cyclin D1 / c-Myc / Pygo-1 | 0.839 | 0.0541 | 0.720 to 0.922 | **P < 0.0001** | **77.97%** |
| Cyclin D1 / Fra-1 / Pygo-1 | 0.853 | 0.0527 | 0.736 to 0.932 | **P < 0.0001** | **77.97%** |
| c-Myc / Fra-1 / Pygo-1 | 0.729 | 0.0654 | 0.597 to 0.836 | P = 0.0179 | 66.10% |
| Cyclin D1 / c-Myc / Fra-1 / Pygo-1 | 0.853 | 0.0509 | 0.736 to 0.932 | **P = 0.0001** | **77.97%** |

### Co-localisation analysis of Wnt signaling target proteins in NL, CC and HCC

Comparative co-localisation analysis was performed on high magnification, deconvolved, IF images to measure the spatial overlap of two proteins in NL, CC and HCC (Fig. 5). Pearson correlation coefficients (r_p_) were compared for 6 permutations (in sets of co-localisation of 2 proteins at a time) in 3 conditions (NL vs CC, NL vs HCC and CC vs HCC), thus yielding 18 comparisons (Fig. 6). Significant differences were observed in the co-localisation between: Pygo-1/c-Myc, Pygo-1/Fra-1, Cyclin D1/Fra-1 and c-Myc/Fra-1 in NL vs CC; whereas in NL vs HCC, co-localisation was significantly different for all permutations except Pygo-1/c-Myc. The comparison of CC and HCC revealed significantly different co-localisation for Pygo-1/Cyclin D1, Pygo-1/c-Myc and Pygo-1/Fra-1. These results indicate smaller overlap for Pygo-1/Fra-1 and c-Myc/Fra-1 in NL (as evident by r_p_ values closer to 0) that increases in CC and HCC. Thus, altered co-localisation of both Pygo-1/Fra-1 and c-Myc/Fra-1 could serve as putative biomarkers for the abundant primary liver cancers (CC and HCC, P < 0.0001), with the prior one more promising as it also distinguished CC from HCC (P < 0.05).

**Fig. 5.**
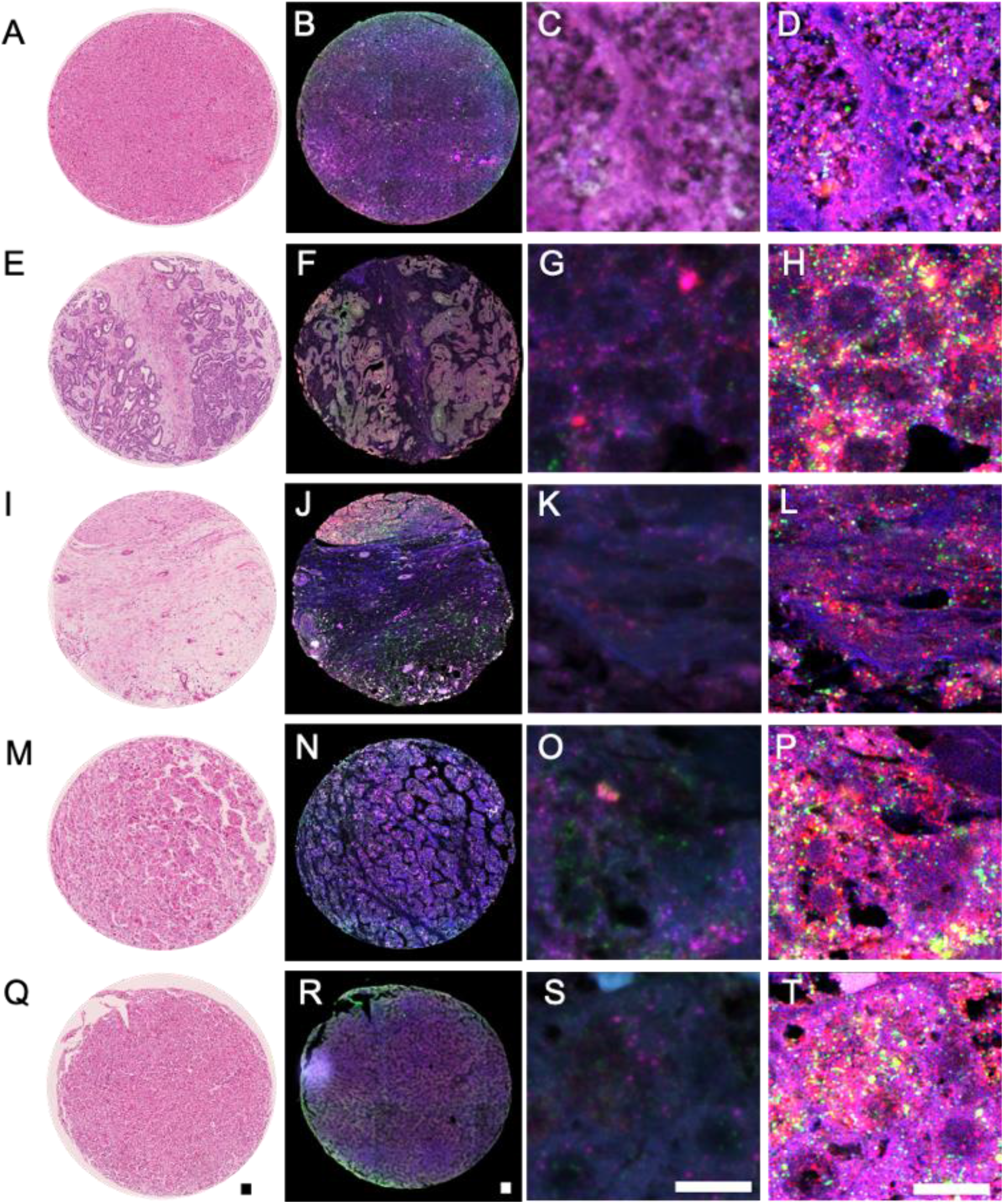
Multi-labeled, high-magnification immunofluorescence for Cyclin D1, c-Myc, Fra-1 and Pygo-1. Representative micrographs for the co-expression of Cyclin D1 (FITC-green), c-Myc (Cy3-red), Fra-1 (Cy5-magenta) and Pygo-1 (Coumarin-blue) in NL (A-D), iCC (E-H), pCC (I-L), pHCC (M-P) and wHCC (Q-T). Whole tissue cores (B, F, J, N, R) were imaged using a Zeiss Axioscan Z.1 slide scanner (Carl Zeiss) at 20x (Supporting Figs S3-S13). For quantitative co-localisation, randomly selected areas from 3 tissue cores each of NL, CC and HCC were imaged (~ 3 areas per core) using Leica SP8 confocal microscope at 63x (NA 1.44) with 6x optical zoom (C, NL; G, iCC; K, pCC; O, pHCC; and S, wHCC). Images were deconvolved (D, NL; H, iCC; L, pCC; P, pHCC; and T, wHCC) using Huygen’s professional software, and co-localisation co-efficients were calculated for further analysis (see Methods). A, E, I, M and Q represent corresponding H&E images for NL, iCC, pCC, pHCC and wHCC respectively. Scale bar = 50 μm for full core images and 10 μm for corresponding high-magnification images.

## Discussion

TAs with a large number of samples, that could be assayed simultaneously, offer an effective use of human samples to investigate the expression of proteins involved in human cancers. We have previously studied the quantitative expression of Wnt-related proteins in several cancers based on multi-labeled qIHC approach using TAs. (18, 19) In this study, we used a similar approach to investigate the role of Wnt signaling across several types of liver tumors to identify putative prognostic and/or diagnostic biomarkers involved in the disease progression.

Wnt signaling, a cell fate developmental pathway, is important for liver regeneration. (20, 21) During liver injury, Wnt proteins initiate liver regeneration by triggering Wnt-β-catenin mediated gene transcription. β-catenin translocation occurs within minutes of partial hepatectomy and leads to the upregulation of c-Myc and Cyclin D1, thus promoting cell proliferation. (20, 21) The current study was designed to investigate whether known downstream target proteins (Pygo-1, c-Myc, Cyclin D1 and Fra-1) of the Wnt signaling pathway are dysregulated across several types of liver tumors in humans. This type of study is important to unravel the molecular events involved in human hepatic carcinogenesis that may directly reflect *in vivo* changes occurring in liver tumors.

Pygo-1, a hydrophilic and non-transmembrane protein, (22) is a part of the nuclear TCF/LEF/β-catenin/Legless/Pygo complex, and acts as a coactivator to regulate the Wnt/β-catenin-mediated transcription of downstream targets. (13) In this study, we observed significantly reduced expression of Pygo-1 in pHCC, iCC, HB, FNH and CRC-LM, vs NL. The combined subtypes of CC and HCC revealed reduced expression of Pygo-1 in both CC and HCC vs NL, which implies its significance to distinguish these cancers from NL. To our knowledge, this is the first study to investigate the expression of Pygo-1 in liver tumors.

c-Myc, a basic helix-loop-helix/leucine zipper transcription factor, (23) heterodimerizes with Max, another leucine zipper protein, and subsequently binds to E-box sequences to activate transcription. (24) Activated transcription is reversed by the binding of heterodimerized Max and Mnt (an antagonist of c-Myc) to the E-box elements. (24) As a downstream target of c-Myc, the expression of Cyclin D1 is increased by a Mnt-Max to c-Myc-Max switch on E-box sequence in the promoter region of Cyclin D1. (24) In our study, reduced expression of c-Myc in iCC and unaltered expression in pCC implies reduced availability of c-Myc to substitute Mnt for activating transcription, thus supporting the reduced expression of Cyclin D1 observed in both subtypes of CC. Although previous studies have reported overexpressed c-Myc in human CC (Supporting Table S5), (25–27) the combined iCC and pCC expression values in our study did not show a significant difference between CC vs NL, suggesting that the reduced expression of c-Myc might be specific to iCC. Based on the dual functionality of c-Myc in cell cycle progression and apoptosis induction, (28) the reduced levels of c-Myc observed in iCC in the current study might indicate the inhibition of c-Myc-dependent apoptosis.

Cyclin D1, an important cell cycle regulator, also regulates gene expression as a co-activator. (29) The nuclear accumulation of β-catenin activates the transcription of Cyclin D1 through TCF/LEF binding sites within the Cyclin D1 promoter. (16) Cyclin D1 is known to drive tumorigenesis in HCC and CC. (30, 31) Overexpression of Cyclin D1 is known to be associated with the progression of iCC and poor prognosis for iCC patients. (29, 32) Tokumoto et al. (25) observed overexpressed Cyclin D1 and c-Myc in 41.7% human iCC samples with significant correlation found only between Cyclin D1 and β-catenin. Previous studies have shown overexpression of Cyclin D1 in CC (25, 29, 32) and HCC (33–35) (Supporting Table S5). However, we observed significantly reduced expression of Cyclin D1 only in iCC, pCC, HB and PAC-LM. Our results suggest a putative role of Cyclin D1 to distinguish CC from both NL and HCC. There has been some controversy regarding the expression of Cyclin D1 in HB. Although a number of reports suggest an overexpression of Cyclin D1 in HB, (36–38) reduced expression was also observed. (39) Another study reported both upregulation and downregulation of Cyclin D1 in 52% and 35% of HB cases, respectively, with no significant difference observed for c-Myc and Fra-1. (40) Cyclin D1 was also identified as a potential prognostic biomarker for mixed epithelial/mesenchymal HB via survival analysis. (36) Previous studies revealed that the elevated expression of Cyclin D1 in HB is associated with increased proliferation and β-catenin staining. (37, 38) Intriguingly, in all these studies only a subset of cancer cases studied was positive for Cyclin D1 (Supporting Table S5), indicating that high Cyclin D1 expression might be associated with the low survival and/or tumor recurrence.

Fra-1, a member of Fos protooncogene family, plays an important role in the development of epithelial tumors. (41) Fra-1 heterodimerizes with the Jun family members to activate target gene transcription, (17) and is known to regulate cell proliferation, transformation and differentiation. (42) Fra-1 has been reported previously as an important prognostic marker for colon cancer and HCC progression. (43, 44) Our observation of elevated expression of Fra-1 in pHCC vs NL may suggest an association with the poorly differentiated cancer. Indeed, the association of overexpressed Fra-1 with poor overall survival in HCC has been reported previously. (44) Among CC subtypes, the expression of Fra-1 was observed to be significantly reduced in pCC. A previous study has reported an increased Fra-1 expression in primary sclerosing cholangitis and primary biliary cirrhosis, the known risk factors for the development of CC. (45) This suggests that Fra-1 may exacerbate the fibrotic liver and is may be involved in the initial stages of development of CC. Combined analysis of CC and HCC subtypes revealed significantly different expression of Fra-1 in CC vs HCC, which indicates its potential to distinguish these primary hepatic cancers.

The discrepancy regarding the expression of Wnt-related proteins between this study and previous reports could be attributed to several factors. In previous studies, overexpression in a subset of samples may indicate the complex classification of the tumors and multiple types of tumor cells within a single tumor sample. Most of these studies have focused on the nuclear expression only and measured the expression level via visual scoring considering less than 10% staining as negative (Supporting Table S5). In this study, we have investigated an overall expression of these proteins in all tumor biopsies and evaluated their potential to be used as biomarkers, specifically for CC and HCC. ROC analysis revealed Cyclin D1 and Pygo-1 as potential biomarkers for CC (Fig. 4). A combinatorial analysis of these two proteins via binary regression model further strengthened the predictive power for CC cases (Table 1).

In addition to measuring the expression of the four proteins of interest, we have also conducted co-localisation analysis to elucidate any significant alterations in the co-localisation patterns of Wnt target proteins in normal and malignant liver tissues (CC and HCC). This was based upon the notion that altered topography may occur in disease and could serve as an additional biomarker. (46) Also, direct and indirect protein-protein interactions jointly contribute to form functional protein assemblies that may interplay in cell signaling mechanisms. (47, 48) Thus, quantitative measures of these molecular interactions in the tissue may also serve to improve the existing knowledge of cellular mechanisms. We report, for the first time, significant differences in the co-localisation of proteins between NL, CC and HCC (Fig. 6). For example, Pygo-1 and Fra-1 showed weak co-localisation in NL, which was increased to low-medium level in CC and medium-high in HCC.

**Fig. 6.**
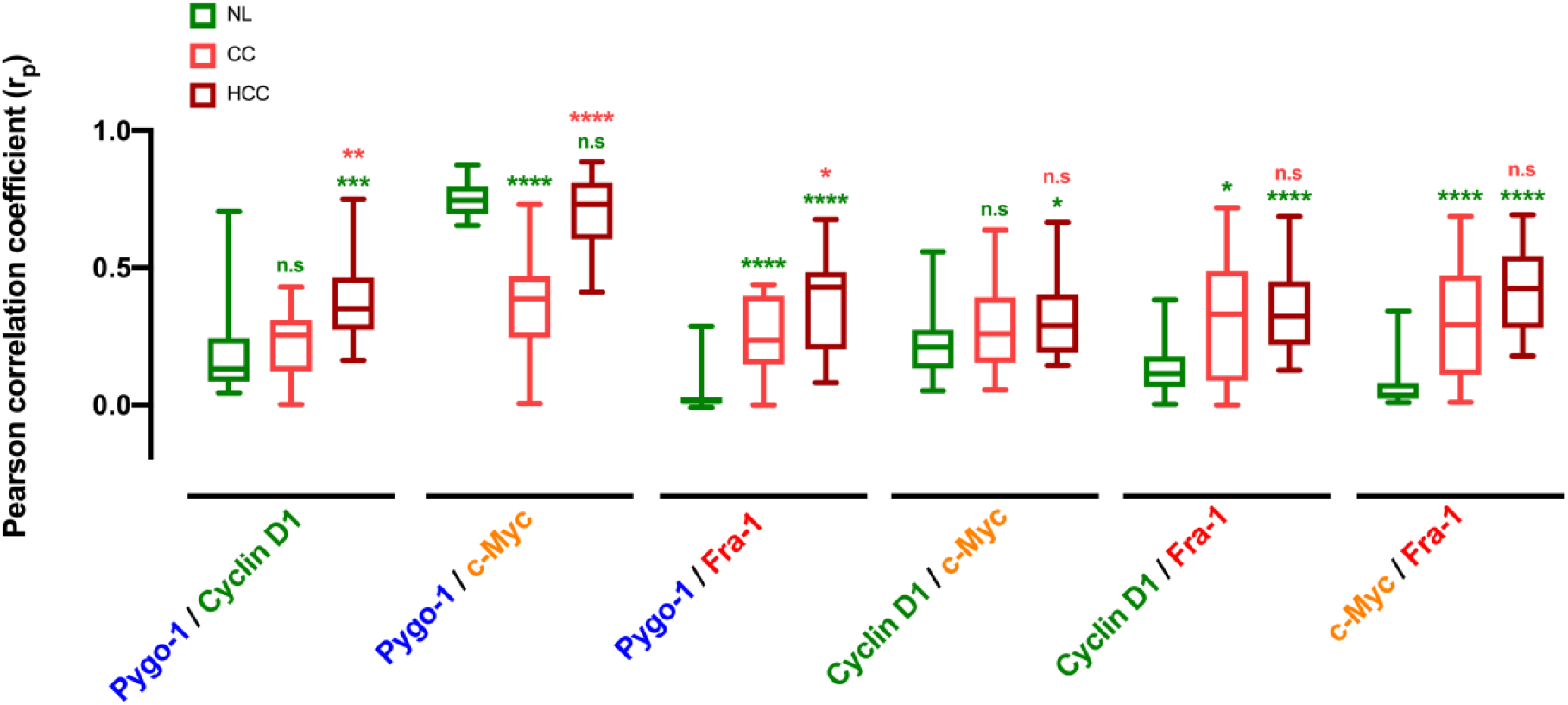
Co-localisation of Wnt signaling target proteins in NL, CC and HCC. Pearson correlation coefficients (r_p_) were calculated for all 6 permutations of the 4 proteins (in the combination of two) from the deconvolved, high magnification immunofluorescent images using Huygens software (n=18). Significance of difference in rp values was calculated using Mann-Whitney U test to distinguish NL vs CC and NL vs HCC, as shown in green above the respective boxes; and CC vs HCC, as shown in pink above the HCC box (* P < 0.05; ** P < 0.01; *** P < 0.001; **** P < 0.0001; and n.s = not significant).

In summary, we characterised the protein expression of four Wnt signaling targets and identified Cyclin D1 and Pygo-1 as putative biomarkers of CC. The comparison of our study with previous findings (Supporting Table S5) indicate the potential of Wnt-related proteins, for the development of therapies targeting the Wnt signaling pathway, as a novel and effective treatment option for liver cancer patients. However, due to the low number of samples in our cohort, analysis of Cyclin D1 and Pygo-1 expression in a larger group of patients enriched for relapse/death would be necessary to validate our findings prior to use these proteins as biomarkers in a clinical trial setting. The expression patterns of Wnt-related proteins may also promote understanding of different mechanisms underlying the pathogenesis of various liver tumors included in this study.

