## Supplementary material for "Comparative analysis of Wnt signaling-related proteins in normal, benign, malignant and metastasised human liver tumors": Table S1

**Table S1.** Histological characterization of human liver tissue samples used to construct human liver tissue microarray.

| Core # | Tissue Type | Sex | Age | Tumour grade/subtype | Tumour stage |
|---|---|---|---|---|---|
| 1 | PAC-LM | F | 82 | Moderately differentiated | |
| 2 | pHCC | M | 57 | Poorly differentiated | |
| 3 | PAC-LM | M | 76 | Moderately differentiated | |
| 4 | CRC Met | M | 73 | Moderately differentiated | |
| 5 | pCC | F | 64 | Adenocarcinoma | T2N0MX |
| 6 | PAC-LM | F | 47 | Well differentiated | |
| 7 | NL | F | 74 | | |
| 8 | iCC | M | 42 | Moderately differentiated | |
| 9 | HCC-CC | F | 68 | Well to moderately differentiated | |
| 10 | FNH | F | 75 | | |
| 11 | HCA | F | 43 | | |
| 12 | pCC | M | 60 | Infiltrating adenocarcinoma | T3N1MXR1 |
| 13 | iCC | M | 62 | Moderately differentiated | |
| 14 | HCA | M | 46 | Well differentiated/ inflammatory | |
| 15 | iCC | F | 59 | Cholangiocellular variant | |
| 16 | FNH | F | 30 | | |
| 17 | HB | M | 1 | HB with focal HCC | |
| 18 | wHCC | M | 61 | Well to moderately differentiated, variable pattern | |
| 19 | PAC-LM | F | 58 | Poorly differentiated adenocarcinoma | |
| 20 | PAC-LM | M | 58 | Moderately differentiated adenocarcinoma | |
| 21 | HCA | F | 53 | Indeterminate subtype | |
| 22 | pCC | M | 73 | Poorly differentiated | pT3N1 |
| 23 | iCC | M | 76 | Well to moderately differentiated | |
| 24 | pCC | F | 65 | Moderately differentiated | pT3N1MX |
| 25 | NL | F | 61 | | |
| 26 | CRC-LM | M | 72 | Moderately differentiated | |
| 27 | pHCC | M | 71 | Poorly differentiated HCC, trabecular and micro-acinar, small cell-type | |
| 28 | HB | M | 3 | Treated HB, wholly epithelial, mixed foetal and embryonal subtype | |
| 29 | wHCC | M | 65 | Well differentiated | |
| 30 | FNH | F | 68 | | |
| 31 | FNH | F | 47 | | |
| 32 | FNH | F | 58 | | |
| 33 | HB | F | 1 | | |
| 34 | FNH | F | 27 | Treated HB (clinical diagnosis), largely matured-appearing. Viable tumour resembling blastema and HCC | |
| 35 | HB | M | 1 | Treated HB with focal residual foetal-type component | |
| 36 | HCA | F | 35 | | |
| 37 | pHCC | M | 75 | Moderately and in places poorly differentiated HCC | |
| 38 | CRC-LM | M | 81 | Moderately differentiated adenocarcinoma | |
| 39 | CRC-LM | M | 65 | Extensively necrotic moderately differentiated adenocarcinoma | |
| 40 | CRC-LM | F | 64 | Intestinal type adenocarcinoma | |
| 41 | NL | M | 70 | | |
| 42 | HCC-CC | M | 63 | Partly necrotic nodule in the right lobe with biphasic differentiation | |
| 43 | HCA | F | 23 | Teleangiectatic (inflammatory) subtype | |
| 44 | iCC | M | 73 | Moderately differentiated adenocarcinoma | |
| 45 | HCC-CC | M | 62 | | |
| 46 | HCA | F | 46 | Telangiectatic / inflammatory subtype | |
| 47 | FNH | M | 49 | | |
| 48 | FNH | F | 29 | | |
| 49 | wHCC | F | 56 | Well to moderately differentiated | |
| 50 | HCA | F | 30 | Well differentiated, inflammatory | |

| | | | | | |
|---|---|---|---|---|---|
| 51 | FNH | F | 21 | | |
| 52 | pCC | M | 29 | Adenocarcinoma | pT4 N1 M0 |
| 53 | pHCC | F | 67 | Poorly differentiated | |
| 54 | FNH | - | 12 | | |
| 55 | HB | M | 2 | Post-chemotherapy, mixed epithelial and mesenchymal type with teratoid features | |
| 56 | HB | F | 2 | Mixed epithelial and mesenchymal type | |
| 57 | wHCC | F | 71 | Well to moderately differentiated | |
| 58 | CRC-LM | M | 72 | Moderately differentiated | |
| 59 | iCC | F | 68 | Adenocarcinoma | pT3aN1Mx |
| 60 | pCC | F | 72 | Moderately differentiated adenocarcinoma | pT2aNxMx |
| 61 | PAC-LM | M | 81 | Moderately to poorly differentiated adenocarcinoma. | |
| 62 | pHCC | M | 82 | Moderately to poorly differentiated, heterogenous histology | |
| 63 | NL | M | 55 | | |
| 64 | HCA | F | 45 | Inflammatory / telangiectatic varian | |
| 65 | HCC-CC | M | 63 | Moderately differentiated | |
| 66 | pCC | F | 64 | Adenocarcinoma | T3 N1 MX |
| 67 | PAC-LM | F | 67 | Moderately differentiated adenocarcinoma | |
| 68 | pCC | F | 74 | Adenocarcinoma | pT3 N1 MX |
| 69 | HB | F | 1 | Mixed epithelial - mesenchymal type | |
| 70 | HB | M | 2 | Mixed type | |
| 71 | wHCC | M | 55 | Well-to-moderately differentiated | |
| 72 | CRC-LM | F | 76 | Moderately differentiated adenocarcinoma | |
| 73 | pHCC | F | 49 | Moderately to poorly differentiated | |
| 74 | HCC-CC | M | 57 | | pT1R0 |
| 75 | pHCC | M | 73 | Multifocal poorly differentiated | |
| 76 | iCC | F | 70 | Moderately differentiated intrahepatic CC | pT2a |
| 77 | pCC | M | 73 | Well differentiated tubular adenocarcinoma | pT4N1Mx |
| 78 | CRC-LM | F | 55 | Moderately differentiate d adenocarcinoma | |
| 79 | wHCC | M | 45 | Well differentiated | |
| 80 | wHCC | M | 71 | Multifocal well differentiated | pT2 |
| 81 | CRC-LM | M | 67 | Moderately differentiated adenocarcinoma | |
| 82 | HCC-CC | M | 58 | Mixed well-differentiated HCC and CC | |
| 83 | pHCC | M | 56 | Moderately and poorly differentiated, multifocal | |
| 84 | pCC | F | 71 | Moderately differentiated | pT3Nx |
| 85 | HCA | M | 59 | Inflammatory sub-type | |
| 86 | pCC | F | 47 | | pT3N1Mx |
| 87 | HCC-CC | F | 59 | | |
| 88 | HCC-CC | M | 60 | | |
| 89 | CRC-LM | F | 75 | Extensively necrotic moderately differentiated adenocarcinoma | |
| 90 | pHCC | F | 18 | Poorly differentiated | |
| 91 | CRC-LM | M | 35 | Moderately differentiated adenocarcinoma | |
| 92 | CRC-LM | M | 68 | Moderately differentiated adenocarcinoma | |
| 93 | HCA | F | 53 | Well-differentiated hepatocellular neoplasm, inflammatory subtype | |
| 94 | HB | F | 2 | Mixed tumour with features of both HB (mixed epithelial and mesenchymal type) and HCC | R1 |
| 95 | pHCC | F | 82 | Poorly differentiated pleomorphic carcinoma | |
| 96 | NL | M | 81 | | |
| 97 | HCC-CC | M | 46 | HCC (moderately differentiated) with foci of cholangiocellular differentiation | |
| 98 | HCC-CC | M | 83 | Poorly differentiated carcinoma with biphasic differentiation | |
| 99 | NL | F | 66 | | |
| 100 | PAC-LM | M | 72 | Moderately differentiated mucinous adenocarcinoma with regressive change | |
| 101 | iCC | M | 67 | Adenocarcinoma- bile duct origin | pT4 NX M1 |
| 102 | HCC-CC | M | 51 | Moderately differentiated | |
| 103 | FNH | F | 67 | | |

| | | | | | |
|---|---|---|---|---|---|
| 104 | HCC-CC | M | 50 | | |
| 105 | FNH | F | 40 | | |
| 106 | PAC-LM | M | 70 | Ductal type moderately differentiated adenocarcinoma | |
| 107 | PAC-LM | F | 30 | | |
| 108 | pHCC | M | 58 | Poorly differentiated | |
| 109 | pHCC | M | 34 | Metastatic HCC, poorly differentiated | |
| 110 | HCC-CC | F | 43 | Mixed malignant hepatobiliary neoplasm with predominantly cholangiocellular features | |
| 111 | HCA | M | 23 | Inflammatory subtype | |
| 112 | NL | M | 75 | | |
| 113 | pHCC | M | 55 | Moderately and focally poorly differentiated HCC | |
| 114 | wHCC | M | 56 | Well differentiated | |
| 115 | wHCC | M | 66 | Well differentiated HCC | |
| 116 | HB | M | 1 | Post treatment HB | |
| 117 | HCC-CC | M | 51 | | |
| 118 | PAC-LM | F | 61 | Moderately differentiated adenocarcinoma | |
| 119 | wHCC | F | 68 | Well to moderately differentiated HCC | |
| 120 | FNH | F | 40 | | |
| 121 | PAC-LM | F | 64 | Adenocarcinoma | |
| 122 | pCC | M | 64 | Moderately differentiated CC | pT2N1Mx |
| 123 | PAC-LM | M | 55 | Moderately differentiated adenocarcinoma | |
| 124 | CR Met | M | 78 | Moderately differentiated adenocarcinoma | |
| 125 | wHCC | M | 53 | Well-moderately differentiated | |
| 126 | iCC | M | 61 | Adenocarcinoma | |
| 127 | pHCC | F | 71 | Poorly differentiated | |
| 128 | pHCC | M | 61 | Poorly differentiated | |
| 129 | NL | F | 11 | | |
| 130 | NL | F | 67 | | |
| 131 | pCC | F | 75 | Moderately differentiated | pT3N1R0 |
| 132 | iCC | F | 44 | | pT3 N0 MX |
| 133 | NL | M | 48 | | |
| 134 | iCC | F | 73 | Moderately differentiated adenocarcinoma | pT2aR1 |
| 135 | pCC | F | 71 | Poorly differentiated adenocarcinoma | pT2N1MX |
| 136 | PAC-LM | M | 55 | Well differentiated adenocarcinoma | |
| 137 | HCA | F | 29 | Indeterminate type | |
| 138 | HB | F | 2 | Treated HB | |
| 139 | CR Met | M | 80 | Moderately differentiated adenocarcinoma | |
| 140 | iCC | F | 45 | Moderately differentiated | |
| 141 | pCC | F | 71 | | pT2 N1 M0 |
| 142 | NL | F | 52 | | |
| 143 | FNH | F | 34 | | |
| 144 | NL | F | 79 | | |
| 145 | wHCC | M | 60 | Well-moderately differentiated, with steatohepatitic-like features | |
| 146 | HCA | F | 40 | Telangiectatic adenoma | |
| 147 | wHCC | M | 58 | Well to moderately differentiated | |
| 148 | FNH | F | 46 | | |
| 149 | NL | F | 75 | | |
| 150 | iCC | F | 47 | Moderately differentiated adenocarcinoma | pT2bN0 |
| 151 | NL | M | 62 | | |
| 152 | wHCC | F | 64 | Well differentiated | |
| 153 | HB | F | 0 | Treated HB with a very focal residual viable epithelial component | |
| 154 | HB | M | 3 | Hepatocellular neoplasm, with features of both HCC and HB | |
| 155 | PAC-LM | M | 84 | Poorly differentiated adenocarcinoma | |
| 156 | HCA | F | 41 | Steatotic hepatocellular adenoma (HNF1-alpha mutation) | |
| 157 | wHCC | M | 57 | Well differentiated | |
| 158 | HB | F | 0 | Post-chemotherapy, mixed epithelial and mesenchymal type | |
| 159 | NL | F | 82 | | |
| 160 | iCC | F | 83 | Peripheral CC | |

| | | | | |
|---|---|---|---|---|
| 161 | iCC | F | 61 | |
| 162 | CRC-LM | M | 68 | Moderately differentiated mucinous adenocarcinoma |
| 163 | HB | M | 0 | High grade, small cell undifferentiated |
| 164 | HCA | F | 24 | Hepatocellular neoplasm (adenoma), with steatosis |
| 165 | HCC-CC | M | 45 | HCC, with features of fibrolamellar variant and combined papillary, cholangiocellular growth pattern |

HuLiv-TMA included 165 tissue cores, with 15 different cores per category of liver tissue. The categories of liver tissue included in the TMA are:

NL: Normal liver tissue (adjacent to colorectal carcinoma metastasis),
pCC: Peri-hilar CC,
iCC: Intrahepatic CC,
pHCC: Poorly differentiated HCC,
wHCC: Well differentiated HCC,
HCC-CC: Combined HCC-CC,
HB: Hepatoblastoma,
FNH: Focal nodular hyperplasia,
HCA: Hepatic adenoma,
PAC-LM: Pancreatic adenocarcinoma liver metastasis, and
CRC-LM: Colorectal carcinoma metastasis samples.
