## Supplementary material for "Comparative analysis of Wnt signaling-related proteins in normal, benign, malignant and metastasised human liver tumors": Table S2

**Table S2.** Quantitative analysis of protein intensities in abundant primary liver cancers.

| Protein | Condition (n) | Percentage Area fraction / Amount of tissue (Mean ± SEM) | Fold change | Significance level | |
|---|---|---|---|---|---|
| | | | | NL vs CC/HCC | CC vs HCC |
| **Cyclin D1** | NL (15) | 38.283 ± 3.503 | | | |
| | CC (29) | 21.821 ± 1.658 | 0.569 | P < 0.0001 | |
| | HCC (30) | 34.717 ± 2.181 | 0.906 | n.s | P < 0.0001 |
| **c-Myc** | NL (15) | 2.200 ± 0.341 | | | |
| | CC (29) | 1.866 ± 0.373 | 0.848 | n.s | |
| | HCC (30) | 1.809 ± 0.308 | 0.822 | n.s | n.s |
| **Fra-1** | NL (15) | 16.683 ± 0.940 | | | |
| | CC (29) | 15.039 ± 1.390 | 0.901 | n.s | |
| | HCC (30) | 24.084 ± 3.132 | 1.443 | n.s | P < 0.01 |
| **Pygo-1** | NL (15) | 1.098 ± 0.146 | | | |
| | CC (29) | 0.690 ± 0.137 | 0.628 | P < 0.001 | |
| | HCC (30) | 0.564 ± 0.094 | 0.514 | P < 0.001 | n.s |

The expression of DAB-stained Wnt target proteins (Cyclin D1, c-Myc, Fra-1 and Pygo-1) was quantified by using an unbiased, semi-automated batch analysis (see Methods). Fold change in protein expression in CC and HCC is relative to NL cores and was confirmed by AUC calculations of data from Fig 3(A). Significance of difference was calculated by Mann-Whitney U test (n.s = not significant).
