## Supplementary material for "Comparative analysis of Wnt signaling-related proteins in normal, benign, malignant and metastasised human liver tumors": Table S3

**Table S3.** Statistical significance of AUC values for ROC curves representing the selectivity and sensitivity of Wnt-related protein targets in different conditions.

| Protein | Comparison (n) | Area under ROC curve | Std. Error | 95% CI | P value |
|---|---|---|---|---|---|
| **Cyclin D1** | NL vs CC (15 + 29) | 0.8552 | 0.05822 | 0.7411 to 0.9693 | **0.0001** |
| | NL vs HCC (15 + 30) | 0.5822 | 0.09584 | 0.3944 to 0.7701 | 0.3730 |
| | CC vs HCC (29 + 30) | 0.8264 | 0.05489 | 0.7188 to 0.9340 | **<0.0001** |
| **c-Myc** | NL vs CC (15 + 29) | 0.6368 | 0.08406 | 0.4720 to 0.8015 | 0.1407 |
| | NL vs HCC (15 + 30) | 0.6044 | 0.08692 | 0.4341 to 0.7748 | 0.2578 |
| | CC vs HCC (29 + 30) | 0.5080 | 0.07711 | 0.3569 to 0.6592 | 0.9155 |
| **Fra-1** | NL vs CC (15 + 29) | 0.6138 | 0.08406 | 0.4720 to 0.8015 | 0.1407 |
| | NL vs HCC (15 + 30) | 0.5644 | 0.08457 | 0.3987 to 0.7302 | 0.4850 |
| | CC vs HCC (29 + 30) | 0.6724 | 0.07003 | 0.5352 to 0.8097 | **0.0229** |
| **Pygo-1** | NL vs CC (15 + 29) | 0.7609 | 0.07429 | 0.6153 to 0.9065 | **0.0050** |
| | NL vs HCC (15 + 30) | 0.7778 | 0.07010 | 0.6404 to 0.9152 | **0.0026** |
| | CC vs HCC (29 + 30) | 0.5230 | 0.07648 | 0.3731 to 0.6729 | 0.7617 |

Quantitative expression values for Cyclin D1, c-Myc, Fra-1 and Pygo-1 were analysed to investigate the diagnostic potential of these proteins for CC and HCC in comparison to NL, and CC vs HCC. AUC values, significance level and 95% confidence interval values for ROC curves distinguishing two conditions are given in the table. Statistically significant values ($P<0.05$) are highlighted in green.
