## Supplementary material for "Comparative analysis of Wnt signaling-related proteins in normal, benign, malignant and metastasised human liver tumors": Table S4

**Table S4.** List of the Antibodies used for DAB and multi-labelled immunofluorescent staining

| Name | Supplier | Cat no. |
|---|---|---|
| Pygo-1 | Novus Biologicals | NBP1-42665 |
| c-Myc | Novacastra/Leica | NCL-c-Myc (Clone 9E11) |
| Fra-1 | Abcam | ab117951 (Clone 12F9) |
| Cyclin D1 | Santa Cruz | sc-718 (Clone M-20) |
| Goat anti Rabbit HRP fab | Abcam | ab102279 |
| Goat anti Mouse HRP fab | Abcam | ab98659 |
