## Supplementary material for "Comparative analysis of Wnt signaling-related proteins in normal, benign, malignant and metastasised human liver tumors": Table S5

**Table S5:** Literature survey of the expression values of the protein targets used in current study with the disease type, experimental method used and representative references.

| Disease type (n) | Protein targets | Expression level (%) | Section type | Experimental method | References |
|---|---|---|---|---|---|
| iCC (27) vs Adjacent normal (9) | Cyclin D1 | ↑ (Semi-quantitative scoring) | TA | qRT-PCR \| IHC \| WB | (Qi, Wang et al. 2019) |
| eCC (18) vs GB (18) | Cyclin D1 | ↑ (44.4%) | TA | IHC \| WB | (Kim, Han et al. 2009) |
| CC (53) vs Normal bile ducts (20) | Cyclin D1 | ↑ (62.3%) | Whole face | IHC | (Zhao, Lu et al. 2008) |
| iCC (42) vs Adjacent normal (42) | Cyclin D1 | ↑ (61.9%) | Whole face | IHC | (Kang, Kim et al. 2002) |
| iCC (66) | Cyclin D1 | ↑ (42%) | Whole face | IHC | (Sugimachi, Aishima et al. 2001) |
| iCC (24) | Cyclin D1<br>c-Myc | ↑ (41.7%)<br>↑ (41.7%) | | IHC \| Sequencing | (Tokumoto, Ikeda et al. 2005) |
| CC (48) | c-Myc | ↑ | Whole face | qRT-PCR \| WB | (Wu, Li et al. 2017) |
| CC (5) | c-Myc | ↑ | Whole face | IHC | (Yang, Liu et al. 2016) |
| HCC (147) vs Adjacent normal (147) | Cyclin D1 | ↑ (41%)<br>(59%) | Whole face | IHC \| IB | (Wu, Lan et al. 2018) |
| HCC (50) | Cyclin D1 | ↑ (58%) | Whole face | IHC | (Joo, Kang et al. 2001) |
| HCC (29) vs Adjacent normal (11) | Cyclin D1 | ↑ (58.6%)<br>(18.2%) | Whole face | IHC | (Yang and Si 2001) |
| HCC (284) vs CLD+NL (174) | c-Myc | ↑ Gene (HCC)<br>↓ Protein (HCC)<br>↑ (CLDs) | TAs | Dual color FISH \| IHC | (Chan, Guan et al. 2004) |
| HCC (66 – IHC + 19 – qRT-PCR \| WB) | Fra-1 | ↑<br>57.6% - tumour<br>21.2% - peritumour | Whole face | IHC \| qRT-PCR \| WB | (Gao, Ge et al. 2017) |

| | | | | | |
|---|---|---|---|---|---|
| HB (104) | Cyclin D1 | ↑ (34%) | TA | IHC | (Purcell, Childs et al. 2012) |
| HB (12) | Cyclin D1<br>c-Myc | ↑ (33%)<br>↑ (83%) | Whole face | IHC | (Ranganathan, Tan et al. 2005) |
| HB (23) vs<br>Adjacent normal (23) | Cyclin D1<br>c-Myc<br>Fra-1 | ↑ (52%)<br>↓ (35%)<br>No significance | Whole face | RT-PCR | (Koch, Waha et al. 2005) |
| HB (14) | Cyclin D1 | ↑ (83%) | Whole face | IHC | (Takayasu, Horie et al. 2001) |
| HB (17) | Cyclin D1 | ↑ (76%) | Whole face | IHC | (Kim, Ham et al. 1998) |
| HB (7 – RNA +<br>4 – protein) | Cyclin D1 | ↓ RNA (57%)<br>↑ Protein (75%) | Whole face | RT-PCR \| IB | (Iolascon, Giordani et al. 1998) |
| HCC vs NL | Fra-1 | ↓ | Whole face | RT-PCR \| IHC | (Kireva, Erhardt et al. 2011) |
| CRC (12 – WB +<br>39 – IHC) | Cyclin D1<br>Fra-1<br>c-Myc | ↑<br>↑<br>↓ | Whole face | WB \| IHC | (Wang, Wang et al. 2002) |

↑ = Increased expression
↓ = Reduced expression
GB = Gall bladder cancer
IB = Immunoblotting
WB = Western blotting
