## Supplementary material for "Comparative analysis of Wnt signaling-related proteins in normal, benign, malignant and metastasised human liver tumors": Fig S1

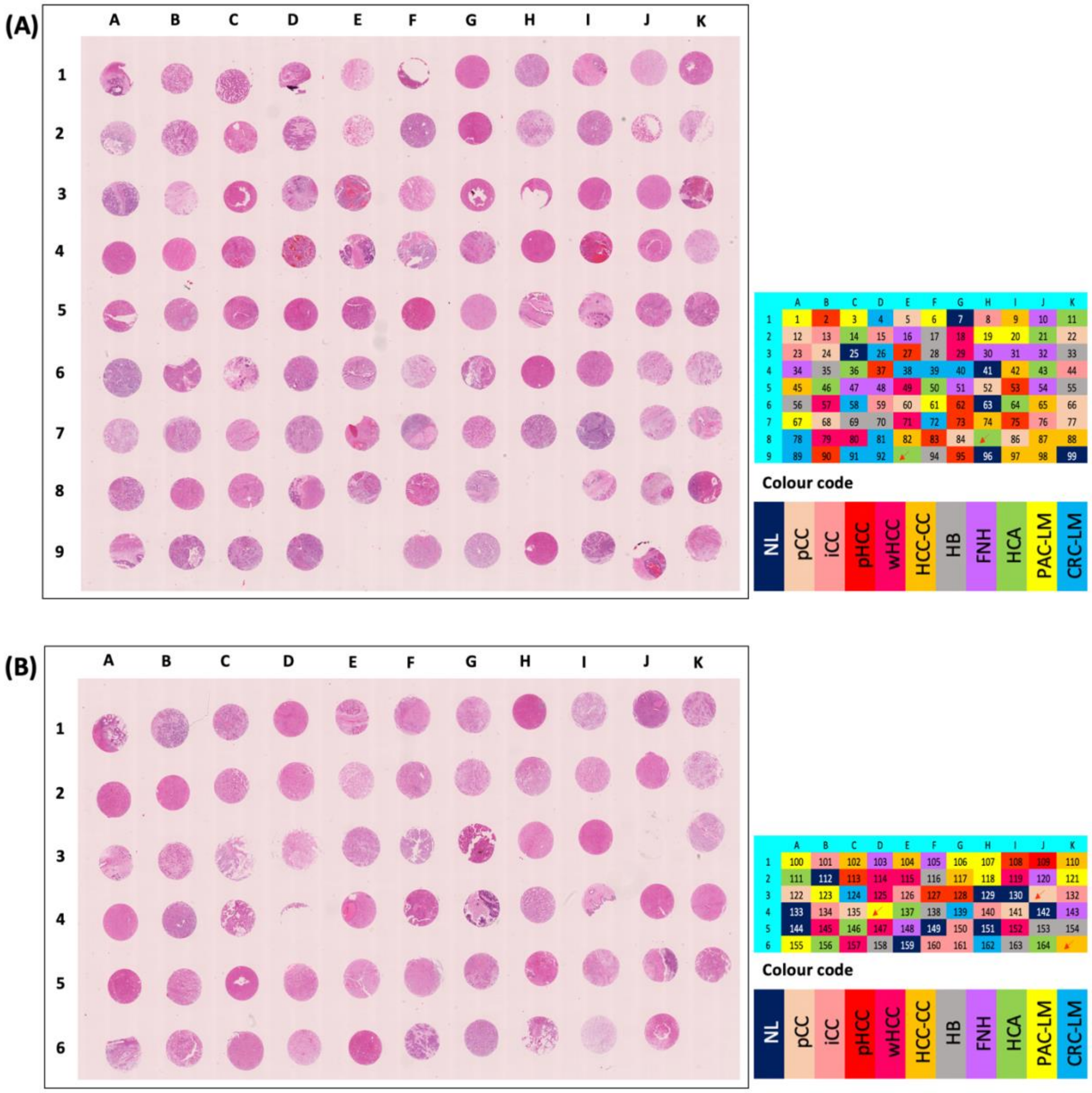


**Fig. S1 Orientation of both blocks of HuLiv-TMA along with the colour code for diseased liver tissue categories.** (A) HuLivTMA-I, consisting of sample cores 1 to 99. (B) HuLivTMA-II, consisting of sample cores 100 to 165. Arrows represent the missing cores. Tissue types include, NL: Normal liver tissue (adjacent to colorectal carcinoma metastasis), pCC: Peri-hilar CC, iCC: Intrahepatic CC, pHCC: Poorly differentiated HCC, wHCC: Well differentiated HCC, HCC-CC: Combined HCC-CC, HB: Hepatoblastoma, FNH: Focal nodular hyperplasia, HCA: Hepatic adenoma, PAC-LM: Pancreatic adenocarcinoma liver metastasis, and CRC-LM: Colorectal carcinoma metastasis.
