## Supplementary material for "Comparative analysis of Wnt signaling-related proteins in normal, benign, malignant and metastasised human liver tumors": Fig S2

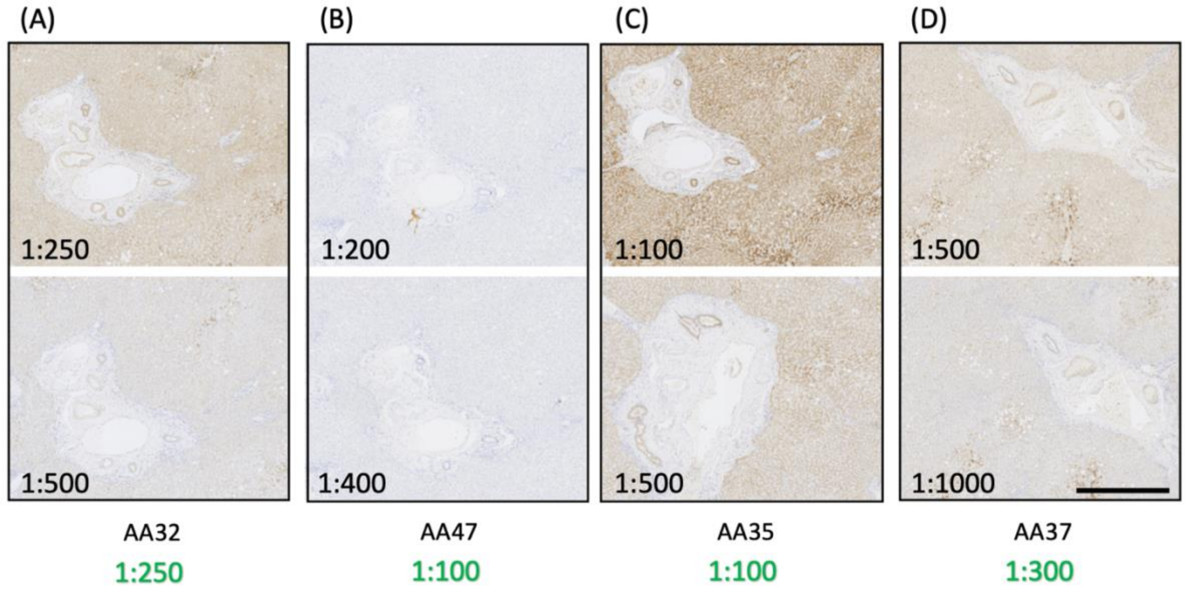


**Fig. S2 Optimization of antibodies for Wnt signalling targets on human liver tissue.** The antibodies were coded (AAxx, A to D) to perform blinded experiments and the concentration of the antibodies was optimized on FFPE whole sections of normal human adult liver (as shown in green), prior to their use in immunostaining of HuLiv-TMA. The slides were imaged at 40x magnification (226 nm/pixel resolution) using a Nanozoomer. AA47 appeared to be completely negative on whole normal liver sections but was used for the HuLiv-TMA staining because of its likeliness to show up in cancer. (Scale bar = 500 μm)
