## Supplementary material for "Comparative analysis of Wnt signaling-related proteins in normal, benign, malignant and metastasised human liver tumors": Fig S5

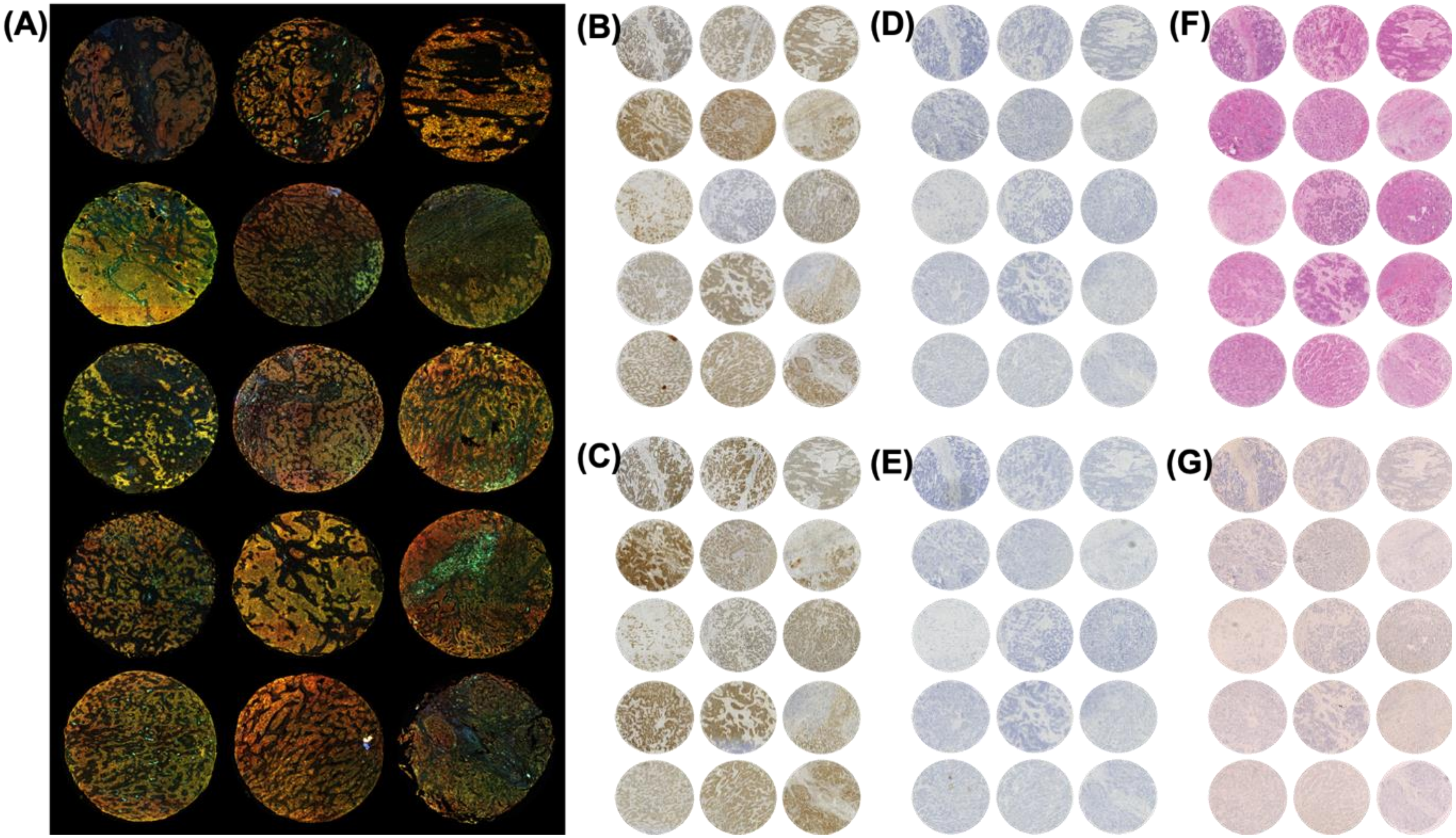


**Fig. S5 Expression of Wnt signalling target proteins in Intrahepatic Cholangiocarcinoma samples.**

(A) Composite images of multiplex immunofluorescence-stained iCC cores (labelled as Cyclin D1-FITC, c-Myc-Cy3, Fra-1-Cy5 and Pygo-1- Coumarin); were acquired using Axioscan (Zeiss) at 20x magnification. (B – E) Singleplex DAB-stained images of iCC cores for individual antibodies, including Cyclin D1, Fra-1, c-Myc and Pygo-1 respectively. H&E-stained images and Negative control images of the corresponding iCC cores are shown in F and G, respectively. All the images for B – G were acquired on Hamamatsu Nanozoomer scanner at 40x magnification.
